# Discordant effects of resource limitation on host survival and systemic pathogen growth in *Drosophila*-bacteria infection models: resistance vs. tolerance

**DOI:** 10.1101/2022.04.29.490073

**Authors:** Aabeer Basu, Aparajita Singh, Suhaas Sehgal, Tanvi Madaan, Nagaraj Guru Prasad

## Abstract

In the experiments presented here, we explore the effect of resource limitation, in form of starvation (which leads to decrease in accessible resources and depletion of reserves) and sexual activity (which leads to reallocation of resources from somatic defence towards reproduction), on immune function of female *Drosophila melanogaster* flies. We infected females with five bacterial pathogens and measured their post-infection survival when subjected to either starvation or sexual activity (mating). Additionally, we measured within host pathogen levels in case of three of these pathogens. Based on previous literature, we predicted that both modes of resource limitation will increase post-infection mortality, but only sexual activity will lead to increase of pathogen load (because of compromised immune function), while starvation will either not affect or reduce pathogen loads (because of reduced availability of resources for the pathogen to proliferate within the host). Our results indicate that both starvation and sexual activity can lead to increased within-host pathogen levels, in addition to increased post-infection mortality, but in a pathogen-specific manner.

## INTRODUCTION

Maintaining a functional immune system and successfully deploying it following a pathogen challenge comes at considerable cost to the host organism in terms of energy and resources [1, 2, 3]. Immune function has thus been canonically considered to be contingent upon hosts access to resources and physiological trade-offs resulting from allocation of available resources among different organismal function [4, 5]. Based on these assumptions suppression of immune system has been predicted to occur under stressful conditions, such as reduced access to nutrition [6]. But recent theoretical developments point towards a picture more complex than simple immune suppression in low resource environments [7].

A resource limited host may not exhibit an increased susceptibility to infections on every occasion because of multiple possible reasons. Even when trade-offs do exist, resource allocation priorities can change depending upon the availability of resources [8]. When under duress it may be advantageous to invest into somatic defence (including immune function), thereby prolonging life-span, rather than investing towards other faculties such as reproduction. Additionally, instead of a global downregulation of the immune system, resource deprivation can induce a restructuring of the immune network to a new stable state that helps maximise immune defence under resource limited conditions [9, 10]. Such restructuring, in principle, can lead to pathogen specific infection outcomes. Furthermore, because the pathogen is dependent on the host for resources for its own proliferation, reducing host access to resources can also affect within host pathogen growth, thereby influencing infection outcome [11]. Reduced uptake of nutrition can often – but not always – be beneficial for infected hosts [12], and infected hosts do also modify their diet to suit immediate energetic requirements [13].

The effect of resource limitation on insect immune function can be dependent on both the severity of limitation (ranging from dilution of nutrition to complete starvation) and on the specific nutritional component missing from the diet [9, 14]. To keep matters simple, we only focused on complete starvation in the experiments presented here. Physiological consequences of starvation in *Drosophila melanogaster* have been studied in great detail from the viewpoint of stress resistance and life-history processes [reviewed in 15, 16], but the effect of starvation on immune function is less well studied. *Relish* deficient flies survive better following infection with *Escherichia coli* and *Erwinia carotovora* (but not when infected with *Enterococcus faecalis*) when subjected to a short period of starvation before infection [17]. Negative effect of starvation on immune function has been reported in other insects [18, 19], but these negative consequences of starvation are specific to only certain components of the immune system and are reversible if the insects are allowed to feed again [19].

Effects of poor nutrition in *D. melanogaster* have primarily been investigated via experiments where the diet of the flies is restricted to low levels of nutrition or a particular ingredient (protein, carbohydrate, etc.) is limiting in the diet. Flies subjected to dietary restriction exhibit increased survival when infected with *Lactococcus lactis* and *Pseudomonas aeruginosa* [20], and *Salmonella typhimurium* [21], but decreased survival when infected with *Listeria monocytogenes* [21]. Defence against *E. faecalis* was unaffected by dietary restriction [21]. Flies on low protein diets have increased susceptibility to infection by *Pseudomonas entomophila* [22] but exhibited reduced mortality when infected with *P. aeruginosa* and *Staphylococcus aureus* [23]. Additionally, flies on a diet with low protein-to-carbohydrate ratio survive better when infected with *Micrococcus luteus* compared to flies on a normal diet [24]. In case of both dietary restriction and low protein diets the effect of experimental manipulation is overtly pathogen specific.

The effect of resource limitation on post-infection fitness of the host may be mediated via its effect on within-host pathogen levels. Since pathogens are dependent on the host for acquiring resources necessary for proliferation, a resource limited host also logically implies a resource limited pathogen [11, 25]. Accordingly, within-host sporulation of the microsporidian parasite *Vavraia culisis* increases with increasing access to nutrition of its host *Aedes aegypti* [26]. In *D. melanogaster*, a high carbohydrate diet similarly increases within-host levels of *Providencia rettgeri* [27]. On the other hand, low protein diets lead to greater pathogen burden for *E. coli* and *Lactococcus lactis* [28], but lower pathogen load in case of *P. aeruginosa* and *S. aureus* [23] in flies. *Relish* deficient flies, when subjected to starvation before infection, carry low levels of pathogen burden when infected with *E. coli* and *E. carotovora* [17]. The effect of limited nutrition on within-host pathogen levels is also apparently mixed, and pathogen specific. Therefore, while reduced access to resources/nutrition can compromise a host’s ability to mount an immune response, it can also compromise the pathogens capacity to proliferate, making the outcome of infection (in terms of host survival) dependent on whether the host immune system or the pathogen’s capacity to proliferate is more affected by lack of resources [11, 25].

Here we only draw from the published literature pertaining to experiments carried out on adult insects. It is fairly possible that the rules governing the effects of resource limitation on immune function differ in larva, nymphs, and adult insects [for example 29, 30]. We have also limited ourselves to studies measuring host survival post-infection and within-host pathogen levels.

Sexual activity (mating) leads to increased susceptibility to bacterial infections in *D. melanogaster* females, and mated females also carry a greater systemic pathogen burden [31, 32, 33, 34]. This post-mating immune-suppression is part of reproduction-immunity trade-off observed in many insects and other invertebrates [35, 36], although not in all insects [37]. Post-mating immune-suppression is driven by reallocation of resources away from somatic defence and towards reproduction, primarily production of eggs. Consequently, females lacking a functioning germline do not exhibit post-mating immune suppression [34]. Different components of the male seminal fluid (sperms and accessory gland proteins, especially sex peptide) play an important role in suppressing female immunity [34]. Sex peptide transferred by males during mating increases synthesis of Juvenile Hormone in females, which reduces a female’s ability to mount an immune response against bacterial infection, leading to greater post-infection mortality [38]. In *D. melanogaster*, Juvenile Hormone dictates investment towards reproduction [39] and also suppresses expression of anti-microbial peptide genes [40]. Mating can also slow down translation, leading to a delay in mounting of an immune response against bacterial pathogens [41]. All together these suggest that mating induced immune suppression in flies can be used as a stand-in model for a resource deprived immune system.

In the present study, we explored how starvation and sexual activity (mating) – individually or in concert – affect post-infection survival of female *D. melanogaster* flies when challenged with five different bacterial pathogens (three Gram-negative bacteria: *Providencia rettgeri, Pseudomonas entomophila, Erwinia c. carotovora*; and two Gram-positive bacteria: *Enterococcus faecalis* and *Staphylococcus succinus*). We also quantified the within-host levels of bacteria following infection to test for the effect of resource limitation on systemic pathogen levels. Based on previously published results, both theoretical and empirical (as described above), we predicted that while both starvation and mating will increase post-infection mortality, mating will increase bacterial levels within host but starvation will reduce the same. Our results suggest that starvation and mating can both compromise post-infection survival of the host and encourage within-host pathogen proliferation, albeit in a pathogen dependent manner.

## RESULTS

### Effect of starvation and sexual activity on post-infection survival of females

In the first experiment, we measured the effect of starvation and sexual activity – individually and in concert – on survival of female *Drosophila melanogaster* when infected with entomopathogenic bacteria. Females were distributed into four treatments: virgin-fed (VF), virgin-starved (VS), mated-fed (MF), and mated-starved (MS); VS, MF, and MS represent the three experimental treatments where host’s access to resources has been limited/manipulated. Females were infected with five different pathogens separately and their post-infection survival was recorded. All females were collected as ‘virgins’ at the time of eclosion of adults, and ‘mated’ females were generated by allowing females to mate with males four hours before infections; all males were removed during the infection process. ‘Fed’ flies were allowed *ad libitum* access to standard diet, while the ‘starved’ flies were hosted on non-nutritive substrate (with ample access to water) from the point of infection onwards. The five pathogens used for infection were *Enterococcus faecalis, Erwinia c. carotovora, Providencia rettgeri, Pseudomonas entomophila*, and *Staphylococcus succinus*. Effect of starvation and sexual activity (mating) on post-infection survival of females was statistically tested using mixed-effect Cox proportional hazards analysis (using virgin-fed flies as the reference; table 1) and pair-wise comparison between treatments was carried out using pair-wise Log-rank tests (table S1). (Please refer to Materials and Methods for a detailed protocol.)

**Table 1.** Output of mixed-effects Cox proportional hazards analysis for effect of different treatments on post-infection survival of females when infected with various pathogens. Hazard ratios are relative to the default level, which is set at 1. The default level for “Treatment” is ‘Virgin, Fed’. Hazard ratio greater than 1 implies reduced survival compared to the default level. (VF: virgin, fed; VS: virgin, starved; MF: mated, fed; MS: mated, starved)

|  | Hazards Ratio | Lower CI (95%) | Upper CI (95%) | Z | p-value | Variance (for random factor) |
| --- | --- | --- | --- | --- | --- | --- |
| *(A) Providencia rettgeri* |  |  |  |  |  |  |
| *Treatment* VS | **4.798763** | **3.172589** | **7.258464** | **7.43** | **1.1e-13** |  |
| *Treatment* MF | **1.810362** | **1.139843** | **2.875318** | **2.51** | **1.2e-02** |  |
| *Treatment* MS | **9.328881** | **6.242455** | **13.941312** | **10.89** | **0.0e+00** |  |
| *Replicate* (Random) |  |  |  |  |  | 0.01109032 |
| *(B) Pseudomonas entomophila* |  |  |  |  |  |  |
| *Treatment* VS | **1.431317** | **1.172323** | **1.747530** | **3.52** | **4.3e-04** |  |
| *Treatment* MF | **1.822423** | **1.490903** | **2.227662** | **5.86** | **4.7e-09** |  |
| *Treatment* MS | **1.449053** | **1.183692** | **1.773903** | **3.59** | **3.3e-04** |  |
| *Replicate* (Random) |  |  |  |  |  | 0.1397181 |
| *(C) Erwinia carotovora carotovora* |  |  |  |  |  |  |
| *Treatment* VS | **1.666889** | **1.323855** | **2.098809** | **4.35** | **1.4e-05** |  |
| *Treatment* MF | **1.872733** | **1.489864** | **2.353992** | **5.38** | **7.6e-08** |  |
| *Treatment* MS | **2.547715** | **2.043663** | **3.176087** | **8.31** | **1.1e-16** |  |
| *Replicate* (Random) |  |  |  |  |  | 0.00672299 |
| *(D) Enterococcus faecalis* |  |  |  |  |  |  |
| *Treatment* VS | 1.218650 | 0.9782164 | 1.518179 | 1.76 | 7.8e-02 |  |
| *Treatment* MF | **1.559359** | **1.2530619** | **1.940527** | **3.98** | **6.8e-05** |  |
| *Treatment* MS | 1.223426 | 0.9750129 | 1.535130 | 1.74 | 8.2e-02 |  |
| *Replicate* (Random) |  |  |  |  |  | 0.005795229 |
| *(E) Staphylococcus succinus* |  |  |  |  |  |  |
| *Treatment* VS | **1.478445** | **1.163285** | **1.878990** | **3.20** | **0.00140** |  |
| *Treatment* MF | **1.571853** | **1.246209** | **1.982590** | **3.82** | **0.00013** |  |
| *Treatment* MS | **1.474480** | **1.160173** | **1.873937** | **3.17** | **0.00150** |  |
| *Replicate* (Random) |  |  |  |  |  | 0.0000157206 |

When infected with *P. rettgeri*, all three resource limited treatments – virgin starved (VS; hazard ratio, 95% confidence interval: 4.799, 3.173-7.258), mated-fed (MF; HR, 95% CI: 1.81, 1.140-2.875), and mated-starved (MS; HR, 95% CI: 9.329, 6.242-13.941) females – exhibited decrease in survival compared to virgin-fed (VF) females (figure 1.a, table 1.a). Females from these three treatments also differed from one another in terms of post-infection survival, with MS females exhibiting greatest mortality, followed by VS and MF (figure 1.f, table S1.a).

**Figure 1.**
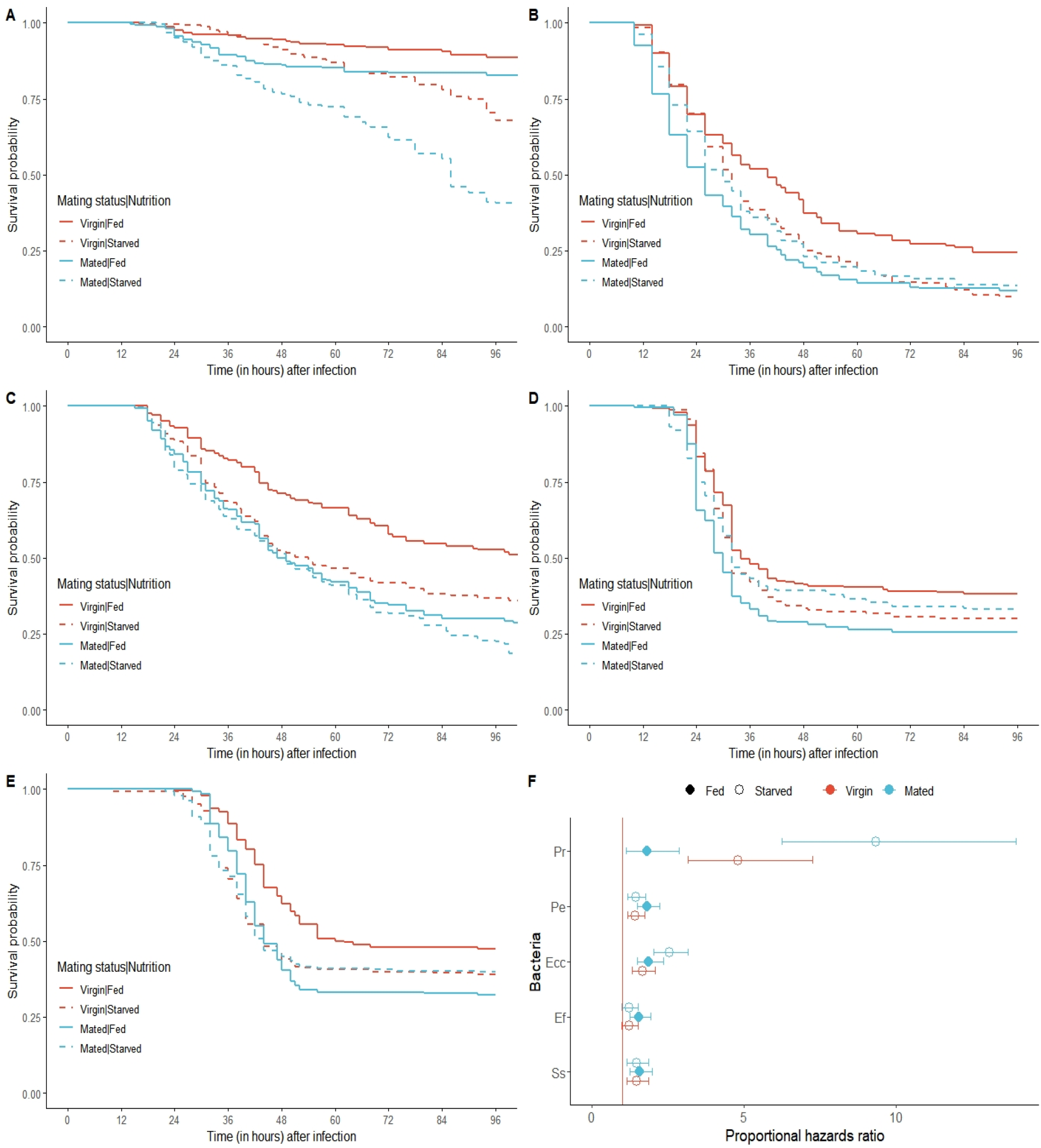
Post-infection survival of females infected with (a) *Providencia rettgeri*, (b) *Pseudomonas entomophila*, (c) *Erwinia c. carotovora*, (d) *Enterococcus faecalis*, and (e) *Staphylococcus aureus* (Survival curves were created after pooling data from all three replicates. Only data from infected females is shown. For a comparison between sham-infected and infected females, refer to supplementary figure S1.); (f) Comparison of hazard ratios (Cox proportional hazards) across different treatments for females infected with each pathogen.

Upon infection with *P. entomophila*, all three resource limited treatments – VS (HR, 95% CI: 1.431, 1.172-1.747), MF (HR, 95% CI: 1.822, 1.491-2.228), and MS (HR, 95% CI: 1.449, 1.184-1.774) females – exhibited decrease in survival compared to VF females (figure 1.b, table 1.b), but these treatments did not differ from one another in terms of post-infection survival (figure 1.f, table S1.b).

Following infection with *E. c. carotovora*, all three resource limited treatments – VS (HR, 95% CI: 1.667, 1.324-2.099), MF (HR, 95% CI: 1.873, 1.490-2.354), and MS (HR, 95% CI: 2.548, 2.044-3.176) females – exhibited decrease in survival compared to VF females (figure 1.c, table 1.c). Additionally, MS females exhibited a significant greater mortality compared to VS and MF females, who did not differ from one another in terms of post-infection mortality (figure 1.f, table S1.c).

Following infection with *E. faecalis*, only MF (HR, 95% CI: 1.559, 1.253-1.940) females exhibited a reduction in post-infection survival compared to VF females; both VS (HR, 95% CI: 1.219, 0.978-1.518) and MS (HR, 95% CI: 1.223, 0.975-1.535) females exhibited survival similar to that of VF females (figure 1.d, table 1.d). Furthermore, there was no significant difference in mortality between MF, VS, and MS females (figure 1.f, table S1.d).

Upon being infected with *S. succinus*, all three resource limited treatments – VS (HR, 95% CI: 1.478, 1.633-1.879), MF (HR, 95% CI: 1.572, 1.246-1.982), and MS (HR, 95% CI: 1.474, 1.160-1.874) females – exhibited decrease in survival compared to VF females (figure 1.e, table 1.e), but these three treatments did not differ from one another in terms of post-infection survival (figure 1.f, table S1.e).

### Effect of starvation and sexual activity on systemic bacterial load in infected females

In the second experiment, using an experimental set-up identical to the first experiment, we measured systemic bacterial load within infected female *Drosophila melanogaster*. Measurement of within-host bacterial load was carried out at two time points – at four and ten hours-post-infection (HPI) – following infection of females separately with three pathogenic bacteria: *Enterococcus faecalis, Providencia rettgeri*, and *Pseudomonas entomophila*. Effect of starvation and sexual activity (mating) on systemic bacterial load of infected females was statistically tested using type III analysis of variance (ANOVA) (table 2) on log-transformed (base 2) bacterial load data, and post-hoc pairwise comparisons were carried out using Tukey’s HSD (tables S2 and S3). (Please refer to Materials and Methods for a detailed protocol.)

**Table 2.**
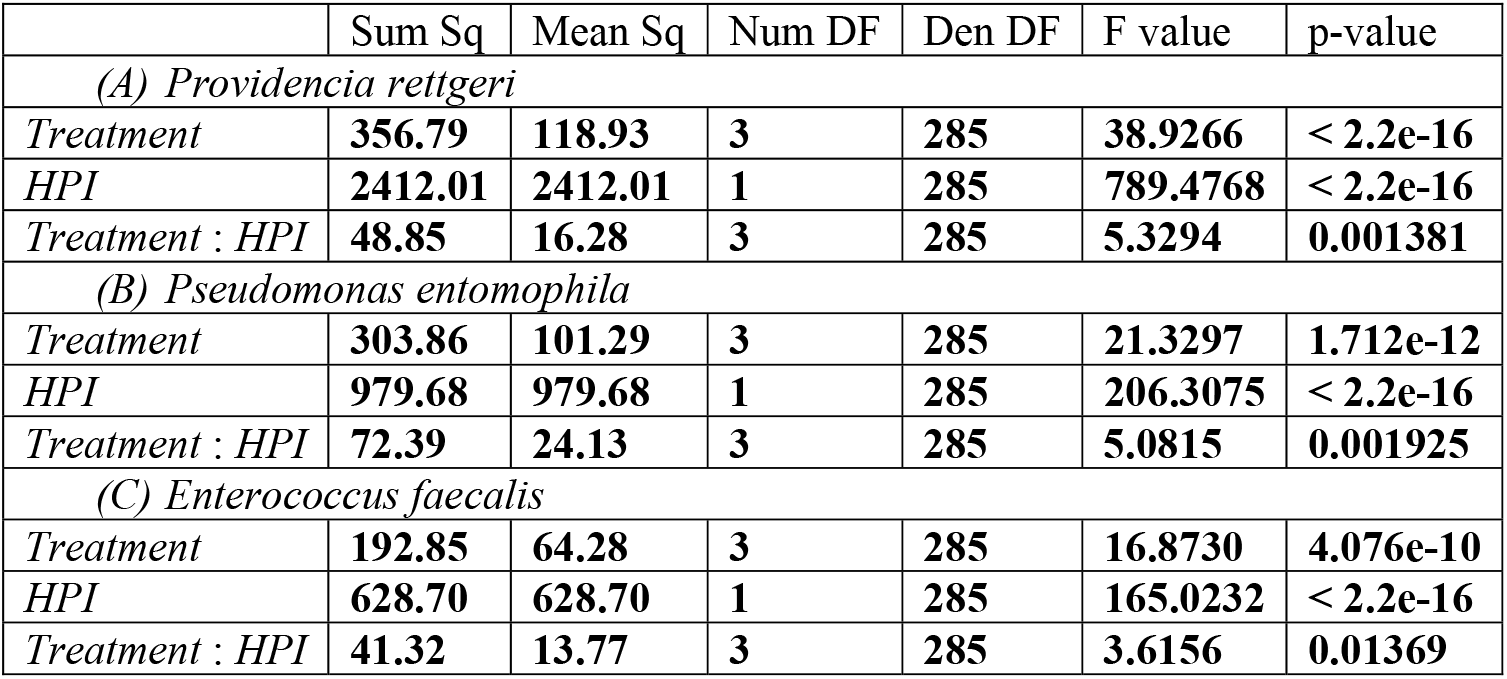
Output of type III analysis of variance (ANOVA) for effect of different treatments and time after infection (HPI) on systemic bacterial load.

|  | Sum Sq | Mean Sq | Num DF | Den DF | F value | p-value |
| --- | --- | --- | --- | --- | --- | --- |
| *(A) Providencia rettgeri* |  |  |  |  |  |  |
| *Treatment* | **356.79** | **118.93** | **3** | **285** | **38.9266** | **< 2.2e-16** |
| *HPI* | **2412.01** | **2412.01** | **1** | **285** | **789.4768** | **< 2.2e-16** |
| *Treatment : HPI* | **48.85** | **16.28** | **3** | **285** | **5.3294** | **0.001381** |
| *(B) Pseudomonas entomophila* |  |  |  |  |  |  |
| *Treatment* | **303.86** | **101.29** | **3** | **285** | **21.3297** | **1.712e-12** |
| *HPI* | **979.68** | **979.68** | **1** | **285** | **206.3075** | **< 2.2e-16** |
| *Treatment : HPI* | **72.39** | **24.13** | **3** | **285** | **5.0815** | **0.001925** |
| *(C) Enterococcus faecalis* |  |  |  |  |  |  |
| *Treatment* | **192.85** | **64.28** | **3** | **285** | **16.8730** | **4.076e-10** |
| *HPI* | **628.70** | **628.70** | **1** | **285** | **165.0232** | **< 2.2e-16** |
| *Treatment : HPI* | **41.32** | **13.77** | **3** | **285** | **3.6156** | **0.01369** |

For females infected with *P. rettgeri*, systemic bacterial load was affected by treatment (F_3,285_: 38.93, p < 2.2e-16), time post-infection (HPI; F_1,285_: 789.48, p < 2.2e-16), and treatment × HPI interaction (F_3,285_: 5.33, p = 0.0014) (figure 2.a, table 2.a). In terms of total bacterial load, VS (LS mean, 95% CI: 12.5, 12.1-13.0), MF (LS mean, 95% CI: 12.3, 11.9-12.8), and MS (LS mean, 95% CI: 14.1, 13.7-14.6) females carried significantly greater bacterial load compared to VF (LS mean, 95% CI: 11.0, 10.6-11.5) females (post-hoc analysis using Tukey’s HSD; table S2.a). MS females also carried a significantly greater load compared to MF and VS females, which did not differ from one another (post-hoc analysis using Tukey’s HSD; table S2.a). At 4 HPI, only the differences between VF and MS females, and VS and MS females, were significant (post-hoc analysis using Tukey’s HSD; table S3.a); at 10 HPI, all pair-wise differences were statistically significant, except the difference between VS and MF females (post-hoc analysis using Tukey’s HSD; table S3.a). Within each treatment, bacterial load always increased significantly with HPI (post-hoc analysis using Tukey’s HSD; table S3.a).

**Figure 2.**
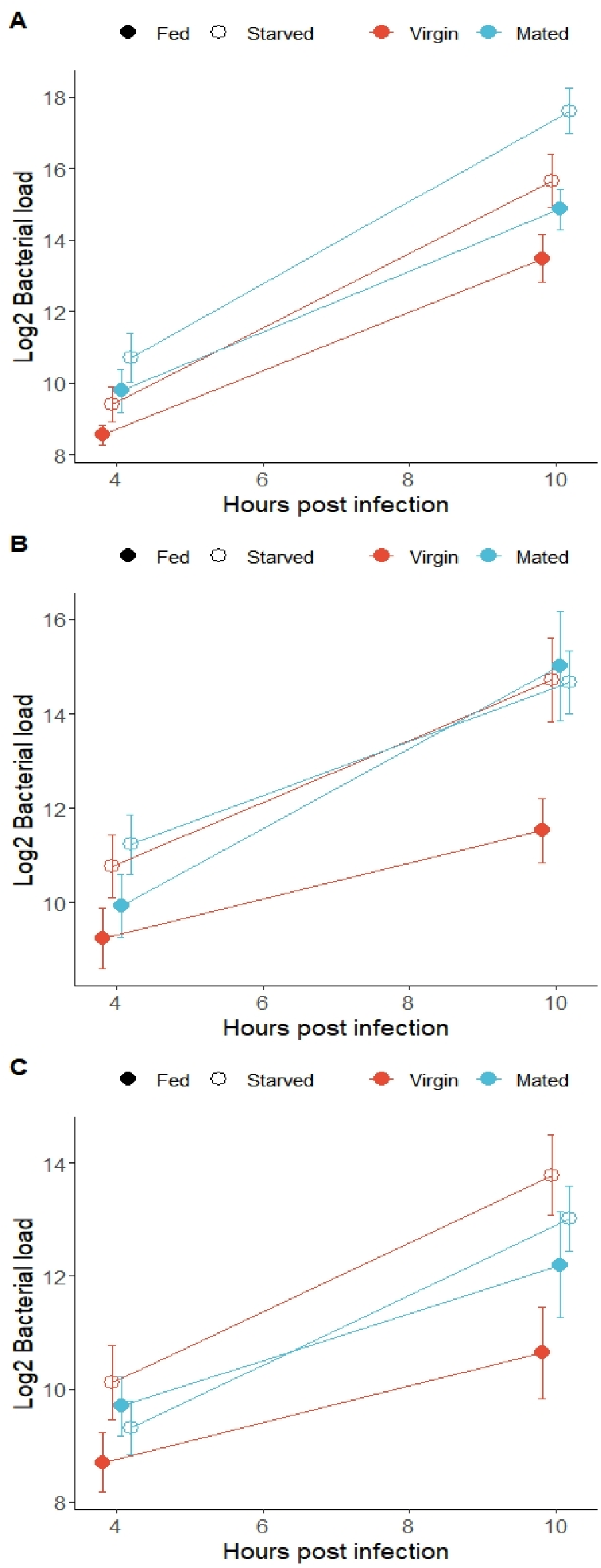
Systemic bacterial load (mean ± 95% confidence intervals) in females infected with (a)*Providencia rettgeri*, (b) *Pseudomonas entomophila*, and (c) *Enterococcus faecalis*. Data from all three replicates are pooled together. For a representation of the full distribution of the data points, refer to supplementary figure S2.

For females infected with *P. entomophila*, systemic bacterial load was affected by treatment (F_3,285_: 21.33, p = 1.71e-12), time post-infection (HPI; F_1,285_: 206.31, p < 2.2e-16), and treatment × HPI interaction (F_3,285_: 5.08, p = 0.0019) (figure 2.b, table 2.b). In terms of total bacterial load, VS (LS mean, 95% CI: 12.7, 11.75-13.7), MF (LS mean, 95% CI: 12.5, 11.48-13.5), and MS (LS mean, 95% CI: 13.0, 11.96-13.9) females carried significantly greater bacterial load compared to VF (LS mean, 95% CI: 10.4, 9.39-11.4) females (post-hoc analysis using Tukey’s HSD; table S2.b); the pair-wise differences between VS, MF, and MS females were not statistically significant. At 4 HPI, only the difference between VF and MS females was significant (post-hoc analysis using Tukey’s HSD; table S3.b); at 10 HPI, VS, MF, and MS females had similar levels of systemic bacteria, and all three had greater bacterial load compared to VF females (post-hoc analysis using Tukey’s HSD; table S3.b). Within each treatment, bacterial load always increased significantly with HPI (post-hoc analysis using Tukey’s HSD; table S3.b).

For females infected with *E. faecalis*, systemic bacterial load was affected by treatment (F_3,285_: 16.87, p = 4.08e-10), time post-infection (HPI; F_1,285_: 165.02, p < 2.2e-16), and treatment × HPI interaction (F_3,285_: 3.62, p = 0.01369) (figure 2.c, table 2.a). In terms of total bacterial load, VS (LS mean, 95% CI: 11.95, 11.42-12.5), MF (LS mean, 95% CI: 10.95, 10.42-11.5), and MS (LS mean, 95% CI: 11.16, 10.64-11.7) females carried significantly greater bacterial load compared to VF (LS mean, 95% CI: 9.67, 9.14-10.2) females (post-hoc analysis using Tukey’s HSD; table S2.c). VS females also had significantly greater bacterial load compared to MF females, but the difference between VS and MS females, and MF and MS females was not statistically significant (post-hoc analysis using Tukey’s HSD; table S2.c). At 4 HPI, females from all four treatments carried similar bacterial load (post-hoc analysis using Tukey’s HSD; table S3.c); at 10 HPI, all pair-wise differences were statistically significant, except the differences between VS and MS females, and MF and MS females (post-hoc analysis using Tukey’s HSD; table S3.c). Within each treatment, bacterial load always increased significantly with HPI (post-hoc analysis using Tukey’s HSD; table S3.c).

## DISCUSSION

In the present study we tested for the effect of starvation and sexual activity (mating), individually or in concert, on post-infection survival and within-host bacterial load of *Drosophila melanogaster* females when infected with bacterial pathogens. The *D. melanogaster* females were divided into four experimental treatments: (a) virgin, fed (VF): *ad libitum* access to resources and no mating; (b) virgin, starved (VS): no access to resources and no mating; (c) mated, fed (MF): *ad libitum* access to resources with mating; and, (d) mated, starved (MS): no access to resources, but with mating (Please refer to Materials and Methods for a detailed protocol). The females were thereafter infected with five bacterial pathogens – *Providencia rettgeri, Pseudomonas entomophila, Erwinia c. carotovora, Enterococcus faecalis*, and *Staphylococcus succinus* – and their post-infection survival was recorded in the first experiment. In the second experiment we measured the within-host bacterial load in infected females, but only for three of the bacterial pathogens: *P. rettgeri, P. entomophila*, and *E. faecalis*.

Among the flies with *ad libitum* access to resources (comparison between VF and MF females), sexual activity increased post-infection mortality for all of the pathogens used for infection, but the difference in mortality between the virgin and mated females was pathogen specific (figure 1.f). Pathogen-specific effect of sexual activity on female immune function has been recorded in previous studies [32]. In case of the three pathogens for which systemic pathogen load was measured, the mated females also carried greater systemic pathogen load compared to virgin females (figure 2.a-c). There is a congruency in results from the two experiments, with increased bacterial levels in the system explaining the increased mortality of mated females. Therefore, sexual activity leads to reduced host resistance, i.e., the ability of the host to control systemic pathogen proliferation (*sensu* Raberg et al [42]), leading to reduced post-infection survival. Mated females carrying greater pathogen load has also been reported in previous studies using *D. melanogaster* [31, 32, 34, 38].

Decrease in host resistance following sexual activity in *D. melanogaster* females may be due to multiple possible reasons. One, mating leads to increased egg production, and depletion of the female fat body to provide resources for the same. Fat body is the primary site for production of anti-microbial proteins required for defence against systemic bacterial infections. Anti-microbial proteins are the primary defence against a few of the pathogens tested here, such as, *P. rettgeri* [43], *P. entomophila* and *E. c. carotovora* [44]. Depletion of fat body can therefore lead to decreased resistance to bacterial infections. Two, mating has been shown to slow down translation of genes in fat body cells, leading to a reduced expression of antimicrobial proteins and other genes relevant for defence against bacterial pathogens, especially *P. rettgeri* [41]. Three, mating can lead to reduced immune-surveillance, because of the dual role of proteins involved in lipid transport: when bound to lipid molecules, proteins such as apolipophorins cannot perform their designated role in pathogen recognition [45, 46]. Four, transfer of lipid reserves from the fat body to ovaries also implies increase in circulating resources, readily accessible to the bacterial pathogen, which can help the pathogen proliferate faster in the body of a mated female [47].

Among virgin females (comparison between VF and VS females), starvation increases mortality in case of four out of five pathogens used for infection (figure 1.f), *E. faecalis* being the exception. Similar to the effect of sexual activity, the degree of change in post-infection mortality of females brought about by starvation was pathogen specific. Also, for all three pathogens for which systemic bacterial loads were measured, VS females carried greater bacterial load compared to VF females (figure 2.a-c), indicating a reduction of host resistance (*sensu* Raberg et al [42]). Resource limitation can lead to restructuring of the immune system [9, 10], which may lead to the effects of resource limitation on insect immune function being pathogen specific. This can explain the pathogen specific results in the experiments presented here. Additionally, the constitutive components of the insect immune system may be less affected by limitation of nutrition compared to the inducible components of the immune system [7]. Phagocytosis, a primarily constitutive defence mechanism, is important for defence against *E. faecalis* [48] in *D. melanogaster*, while defence against *P. rettgeri* is primarily mediated by diptericin [43], an inducible antimicrobial peptide. This might explain why survival following infection with *E. faecalis* is unaffected by starvation, while starved females die more when infected with *P. rettgeri*. This line of reasoning can be extended to include the other two Gram-negative bacteria used in these experiments, *P. entomophila* and *E. c. carotovora*, since defence against these two bacteria is also mediated by inducible anti-microbial peptides in *D. melanogaster* [44]. Anorexia, induced by either dietary restriction or gene mutations that affect feeding behaviour in flies, has also been shown to not affect survival of flies when infected with *E. faecalis*, while survival post-infection with other bacterial pathogens can be compromised [21].

Interestingly, VS females carried significantly greater pathogen load compared to VF females when infected in *E. faecalis* (figure 2.c), although there was no significant difference in mortality between these two treatments (figures 1.d and 1.f). Differences in post-infection survival can be caused by change in host resistance or host tolerance (i.e., the ability of the host to deal with infection induced damage to the soma; *sensu* Raberg et al [42]). Single gene mutations in *D. melanogaster* can affect resistance and tolerance to bacterial infections independent of one another [49], and manipulation of diet can have similar discordant effects on host resistance and tolerance in crickets [50] and mice [51]. We propose that starvation leads to increased tolerance to bacterial infections in flies, especially in case of *E. faecalis* infections. Production of reactive oxygen species is part of the *D. melanogaster* defence repertoire against various bacterial pathogens [52], especially *E. faecalis* [53], which can lead to immunopathology (damage to the host tissue caused by the host immune system; *sensu* Sadd and Siva-Jothy [54], Pursall and Rolf [55]). Starvation is known to induce increased production of antioxidants in Carob moth larvae [56]. Therefore, it is possible that in *D. melanogaster* too, starvation might protect flies from harmful effects of reactive oxygen species, leading to increased tolerance and decreased immunopathology, culminating into improved post-infection survival even with high systemic pathogen loads. The possibility of disease tolerance being contingent upon availability of resources has also been suggested by various previous studies (reviewed in [57]).

There is a disagreement between results from post-infection survival assay and systemic pathogen load measurements in a few other instances. For example, VS females die more compared to MF females when infected with *P. rettgeri* even though they carry similar bacterial loads, and VS females die equally as that of MF females when infected with *E. faecalis* even though in this case VS females carry significantly greater pathogen load. Extending our previous argument, these observations can be explained if starvation had differential effects on resistance and tolerance to different pathogens used in these experiments.

An alternative to increase in host tolerance can be reduction in pathogen virulence. Lack of host nutrition has been shown to increase virulence in viral pathogens of vertebrates [58]. Virulence of a pathogen is a function of its ability to grow within the host (proliferation) and capacity to damage the host (pathogenicity; [59]). Our results (figure 2) clearly show that pathogen proliferation increases when hosts are starved. There is no evidence in literature yet that pathogenicity of a bacteria can be influenced by environmental factors experienced by the host. Therefore, *for the present discussion*, we discount the possibility that host-starvation induced increase in bacterial virulence explains our observations.

A meta-analysis by Pike and colleagues [11] revealed that increasing hosts’ access to nutrition increases within host pathogen fitness in invertebrates and insects, suggesting that increased host nutrition fuels greater within-host pathogen proliferation. Previous results from studies on *D. melanogaster* show that the effect of manipulating host nutrition can both increase and decrease within-host pathogen levels depending upon pathogen identity [17, 23, 27, 28]. Results from our experiments show that for all three pathogens for which bacterial load measurements were made, starved females carried a greater pathogen load compared to females fed ad libitum (comparison of VS and MS females with VF females). Pathogen’s access to resources for proliferation is limited by host’s access to nutrition, but here we observe that starved hosts carry a greater bacterial load. This further suggests that the observed results are driven by starvation-induced reduction in host resistance, and additionally, for at least these three pathogens, limiting host resources does not have a negative effect on the pathogen’s capacity to proliferate. Our results align well with the predictions of Cressler and colleagues [25], who predicted that pathogens have a competitive advantage at low and intermediate resource levels, while the host immune system benefits when resources are abundant.

Lastly, our results suggest that although starvation and mating individually do lead to increased post-infection mortality and higher systemic bacterial loads, subjecting females to a dual treatment of starvation and mating does not significantly alter their immune phenotypes: in case of all but two pathogens (*P. rettgeri* and *E. c. carotovora*), MS females have similar survival and bacterial load as that of VS and MF females (figure 1.f and 2.a-c; tables S1 and S2). We propose that this is driven by the fact that *D. melanogaster* females stop egg production when subjected to starvation [60, 61], making MS females physiologically similar to VS females. Previous studies have shown that *D. melanogaster* females lacking a germline do not exhibit post-mating immune suppression [34; but see 31], and such females also respond differently in terms of gene expression patterns (compared to females with a functioning germline) when subjected to either mating or immune challenge [62, 63]. Starvation induced suspension of reproduction can therefore alleviate the negative effect of mating on female immune function, but in a pathogen specific manner.

To summarize, in the present study we focused on how different modes of resource limitation − starvation (global unavailability of resources) and sexual activity (reallocation of resources away from somatic defence and into reproduction) – affect infection outcome (survival) and within-host pathogen levels in *Drosophila melanogaster* females. Results show that mated (sexually active) females have reduced resistance to bacterial infection, which manifests as increased post-infection mortality. Starvation can also lead to reduced resistance, but conditional upon the mating status of the female fly. Additionally, starvation can increase tolerance to bacterial infection, in a pathogen dependent manner. Therefore, our results suggest that the lack of resources to the immune system, whether it is because of unavailability of nutrition or because of reallocation of resources away from the immune system, can compromise host’s ability to resist systemic pathogen proliferation, but the ultimate infection outcome also depends upon the change in host’s tolerance to infection brought about by resource limitation.

## MATERIALS AND METHODS

### Fly populations and general handling

Experiments reported here were carried out on flies from a large, outbred laboratory adapted population of *Drosophila melanogaster*, LH [64, 65, 66]. The LH population is maintained on a 14-day discrete generation cycle, at 25 °C temperature and 12:12 hour light-dark cycle, on cornmeal-molasses-yeast medium, at a census size of about 1900 adults. The flies are maintained in vials (95 mm height and 25 mm diameter); each generation starts with setting up of 60 vials with 150 eggs each on 8-10 ml of food medium. 12 days post-egg laying (PEL), by which time most adults have eclosed, adults from different vials are mixed together and redistributed into 60 vials with 16 females and 16 males in each vial. The vials are supplied with limiting live dietary yeast supplement, and on 14^th^ day PEL, the adults are transferred to fresh vials and allowed to oviposit for 18 hours to start the next generation. Vials with eggs greater than 150 undergo egg-culling to maintain the specified egg density in vials and avoid crowding.

### Derivation of experimental flies

12-day PEL adults were transferred to plexiglass cages (14 cm length × 16 cm width × 13 cm height) at a density of 1000-1200 flies, and the cages are provided with standard food medium in 60 mm Petri plates. For collection of eggs for setting up experiments, cages are provided with a fresh food plate, supplemented with *ad libitum* live yeast supplement, for 48 hours. This is done to encourage egg production and laying in the females. Following this, a fresh food plate is provided to the cages and 12-14 hours later eggs are collected off these plates (using moist paint brushes on 1.5% agar gel) and seeded into food vials (with 8-10 ml of food medium) at an exact density of 150 eggs per vial. The number of vials set up in this manner depends upon the requirement for a particular experiment. These vials are then incubated under standard conditions (detailed above) for egg to mature into larvae and then into adults. On 10^th^ day PEL, during the eclosion peak, adults are collected as virgins within 5-6 hours of eclosion, and housed in single-sex vials (each with 1-2 ml of food medium) at constant density of 8 females per vial or 10 males per vial. Flies are housed in these vials till further manipulation/experimentation.

### Bacterial handling and infection protocol

The bacterial isolates are preserved as glycerol stocks at -80 °C. To obtain live bacterial cells for infections, 10 ml lysogeny broth (Luria Bertani Broth, Miler, HiMedia) is inoculated with glycerol stocks of the necessary bacterium, and incubated overnight with aeration (150 rpm shaker incubator) at suitable temperature. 100 microliters from this primary culture is inoculated into 10 ml fresh lysogeny broth and incubated for the necessary amount of time to obtain confluent (OD_600_ = 1.0-1.2) cultures. The bacterial cells are pelleted down using centrifugation and resuspended in sterile MgSO_4_ (10 mM) buffer at optical density (OD_600_) of 1.0. Flies are infected, under light CO_2_ anaesthesia, by pricking them on the dorsolateral side of their thorax with a 0.1 mm Minutien pin (Fine Scientific Tools, USA) dipped in the bacterial suspension. Sham-infections (injury controls) are carried out in the same fashion, except by dipping the pins in sterile MgSO_4_ (10 mM) buffer. Across three experiments reported in this paper, five pathogens in total were used in this study, namely

a. *Enterococcus faecalis* [67], incubation temperature 37 °C;
b. *Erwinia carotovora carotovora*, strain Ecc15 [68], incubation temperature 29 °C;
c. *Providencia rettgeri* [62], incubation temperature 37 °C;
d. *Pseudomonas entomophila*, strain L48 [44, 69], incubation temperature 27 °C; and,
e. *Staphylococcus succinus*, strain PK-1 [70], incubation temperature 37 °C.

### Systemic bacterial load estimation

To measure the systemic bacterial load, infected females are first surface sterilised using 70% ethanol for 1 minute and 30 seconds, twice. Females are then washed in sterile distilled water for 30 seconds and dried using autoclaved tissue paper. Females are then transferred individually to 1.5 ml vials (micro-centrifuge tubes) containing 50 or 75 microliters (depending upon pathogen used for infection) of sterile MgSO_4_ (10 mM) buffer. Females are homogenised in these vials using a motorised pestle for 50-60 seconds. This homogenate is serially diluted (1:10 dilutions) 8 times in sterile MgSO_4_ (10 mM) buffer. 10 microliters from each dilution, and the original homogenate, are spotted onto a lysogeny agar plate (2% agar, Luria Bertani Broth, Miler, HiMedia). The plates are incubated at required temperature for 8-12 hours (depending upon pathogen used for infection), and the number of colony forming units (CFUs) in each dilution is counted. The number of CFUs in the *countable* dilution (30 ≤ CFUs ≥ 300) is multiplied by appropriate dilution factor to obtain the bacterial load for each individual female.

### Experiment 1

*Effect of starvation and sexual activity on post-infection survival of females*. Virgin females and males were obtained following the protocol described above. On day 12 PEL, half of the females were randomly assigned to ‘virgin’ treatment and the rest to ‘mated’ treatment. Females in the ‘mated’ treatment were combined with males in fresh food vials (1-2 ml standard food medium) in groups of 8 females and 10 males per vial, and allowed to mate for 4 hours (it was visually confirmed that each female had mated at least once). Following this, the females were lightly anaesthetised and infected with bacterial pathogens (or sham-infected) following the infection protocol described above; males were discarded. Females from the ‘virgin’ treatment were similarly infected (or sham-infected). Following infections, half of the females from both these treatments were housed in vials with 1-2 ml of standard food medium (‘fed’ treatment), and the remaining were housed in vials with 1-2 ml 2% non-nutritive agar gel (‘starved’ treatment). This produced four experimental treatments:

a. Virgin, Fed (VF): 10 vials of infected flies and 5 vials of sham-infected females, each vial with 8 females;
b. Virgin, Starved (VS): 10 vials of infected flies and 5 vials of sham-infected females, each vial with 8 females;
c. Mated, Fed (MF): 10 vials of infected flies and 5 vials of sham-infected females, each vial with 8 females; and,
d. Mated, Starved (MS): 10 vials of infected flies and 5 vials of sham-infected females, each vial with 8 females.

Note that in this experiment females were subjected to starvation from the time of infection. The vials were monitored for mortality every 4-6 hours, for 96 hours post-infection (HPI); alive flies were shifted to fresh food/agar vials at 48 HPI. This experiment was carried out for five bacterial pathogens – *E. faecalis, E. c. carotovora, P. rettgeri, P. entomophila*, and *S. succinus* and replicated thrice for each pathogen. In each replicate, 320 females were subjected to infection (80 females x 4 treatments) and 160 females were subjected to sham-infection (80 females x 4 treatments).

### Experiment 2

*Effect of starvation and sexual activity on systemic bacterial load in infected females*. Following a protocol identical to that of Experiment 1, females were distributed into four treatments (VF, VS, MF, and MS, as described above) and infected; 100 infected females in each treatment and 30 sham-infected females in each treatment. Following infections, females were housed in plexiglass cages (14 cm length × 16 cm width × 13 cm height), with all females from a particular treatment in a single cage; infected and sham-infected females were housed in separate cages. (Cages of ‘fed’ treatments were supplied with standard food medium in 60 mm Petri plates and cages of ‘starved’ treatments were supplied with 2% non-nutritive agar gel in 60 mm Petri plates.) At 4 and 10 HPI, 12 females were randomly aspirated out of cages for each treatment (for infected flies only) and the systemic bacterial load was measured for individual females following the CFU enumeration protocol described above. This experiment was carried out for three bacterial pathogens – *E. faecalis, P. rettgeri*, and *P. entomophila* – and replicated thrice for each pathogen. In each replicate, systemic bacterial load was measured for 72 individual females (12 females x 2 time-points x 4 treatments).

### Statistical analysis

Post-infection survival data from experiment 1 was analysed using a mixed-effects Cox proportional hazards model, including ‘treatments’ as a fixed factor and ‘replicate’ as a random factor. Since sham-infected females exhibited negligible mortality (figure S1), survival data from only the infected females were included for analysis. Systemic bacterial load data from experiment 2 was analysed using type III analysis of variance (ANOVA) on log (base 2) transformed data, including ‘treatment’, ‘HPI’, and ‘treatment × HPI’ interaction as fixed factors, and ‘replicate’ as random factor. Post-hoc analysis for pairwise comparison was carried out using Tukey’s HSD method. All analyses were carried out using R statistical software (version 4.1.0 [71]), using various functions from the *survival* [72], *coxme* [73], *lmerTest* [74], and *emmeans* [75] packages. Graphs were created using the *ggplot2* [76] and *survminer* [77] packages.

## Acknowledgements

The various bacterial pathogens used in this study were obtained from various labs across the world. We thank Prof. P Cornelis (Vrije Universiteit Brussel, Belgium) for providing us with *Pseudomonas entomophila*, Prof. B Lazzaro (Cornell University, USA) for *Enterococcus faecalis* and *Providencia rettgeri*, Dr. E Sucena (Instituto Gulbenkian Ciencia, Portugal) for *Erwinia c. carotovora*, and Dr. K Singh (IISER Mohali, India) for *Staphylococcus succinus*. We also thank Manas Geeta Arun for maintenance of the LH fly populations used in this study.

## SUPPLEMENTARY FIGURES AND TABLES

**Supplementary figure S1.**
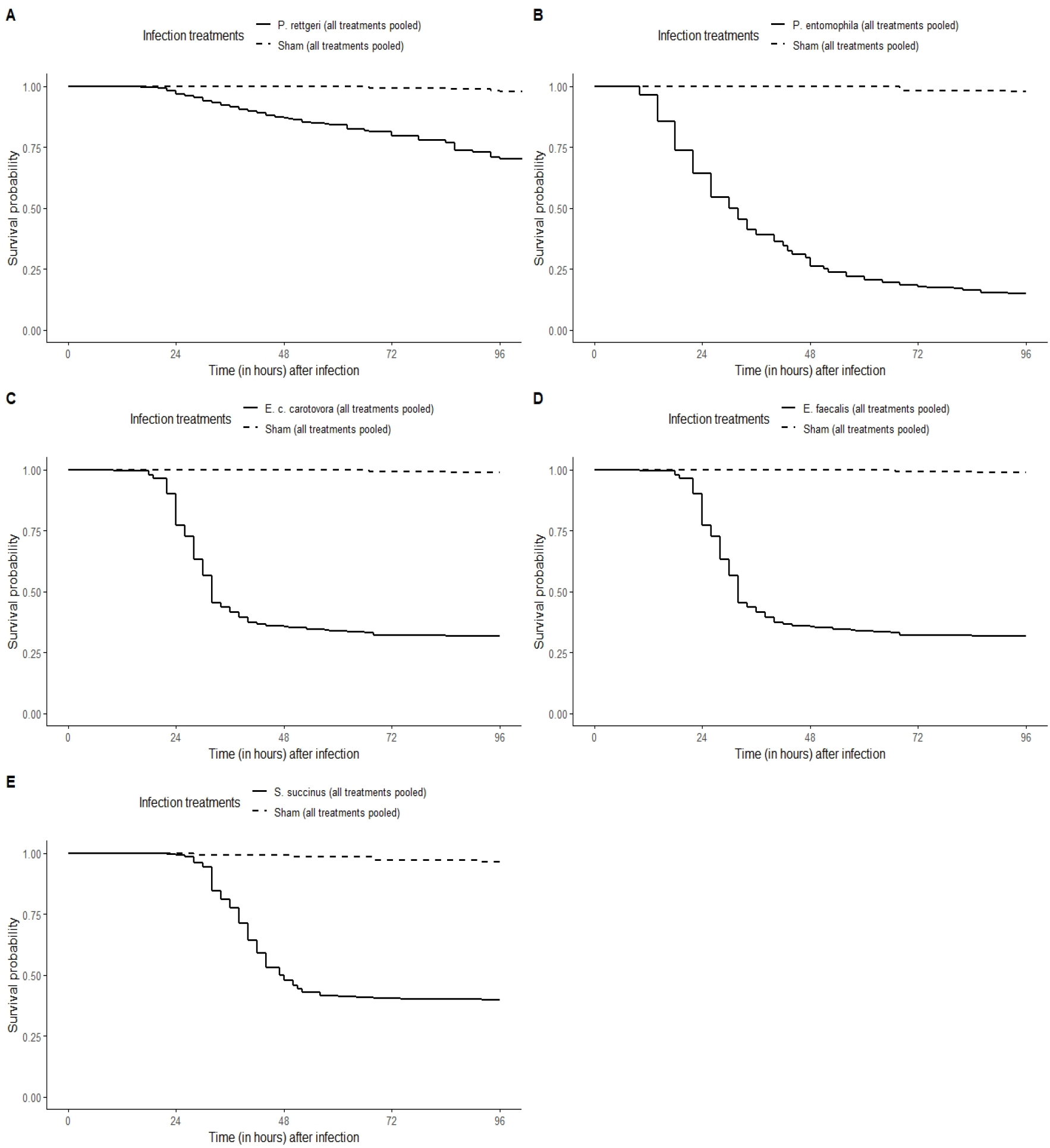
Effect of infection on survival of females. Difference of survival between sham-infected females and females infected with (a) *Providencia rettgeri*, (b) *Pseudomonas entomophila*, (c) *Erwinia c. carotovora*, (d) *Enterococcus faecalis*, and (e) *Staphylococcus aureus*. Survival curves represent females of all treatments and replicates pooled together.

**Supplementary figure S2.**
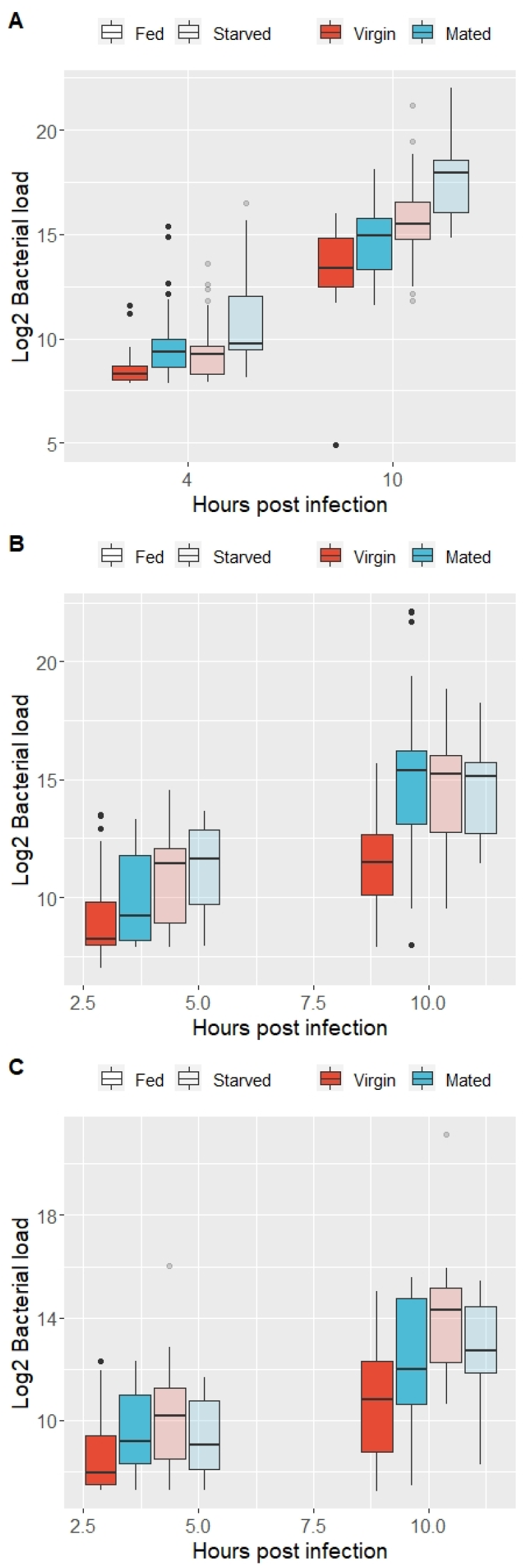
Systemic bacterial load in infected females. Box plots with whiskers (1.5 inter-quartile range) for distribution of systemic bacterial load of individual females (all replicates pooled together) infected with (a) *Providencia rettgeri*, (b) *Pseudomonas entomophila*, and (c) *Enterococcus faecalis*.

**Supplementary table S1.**
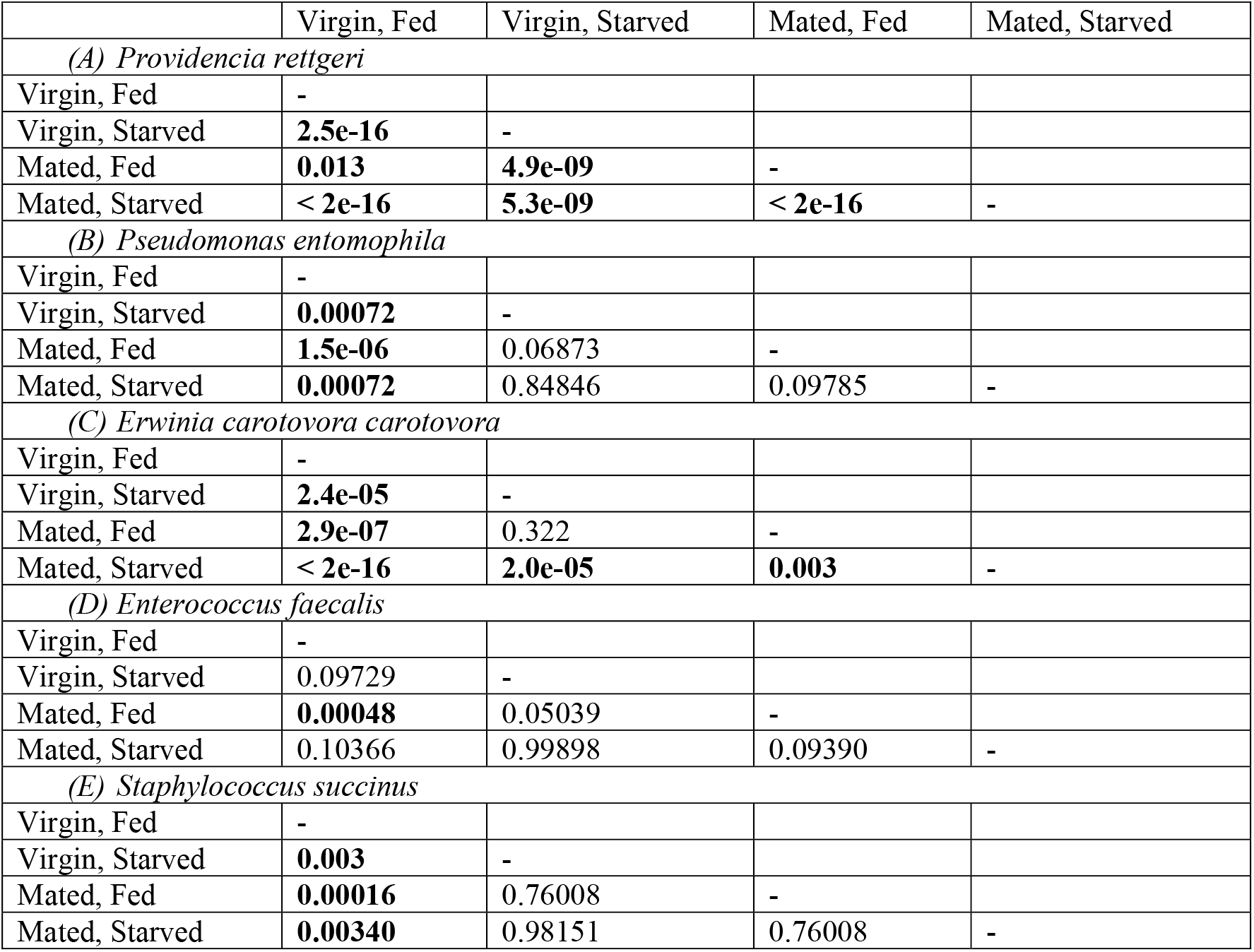
Output of pair-wise Log-rank analysis for effect of different treatments on post-infection survival of females when infected with various pathogens. p-values adjusted for multiple comparisons using Benjamini-Hochberg method.

**Supplementary table S2.** Post-hoc analysis using Tukey’s HSD for pairwise comparisons of the effect of ‘treatments’ on systemic bacterial load in infected females.

| Pairwise comparison | Estimate | SE | DF | t-ratio | p-value |
| --- | --- | --- | --- | --- | --- |
| *(A) Providencia rettgeri* |  |  |  |  |  |
| **Virgin Fed - Mated Fed** | **-1.304** | **0.295** | **292** | **-4.421** | **0.0001** |
| **Virgin Fed - Mated Starved** | **-3.133** | **0.295** | **292** | **-10.622** | **<.0001** |
| **Virgin Fed - Virgin Starved** | **-1.518** | **0.295** | **292** | **-5.146** | **<.0001** |
| **Mated Fed - Mated Starved** | **-1.829** | **0.295** | **292** | **-6.202** | **<.0001** |
| Mated Fed - Virgin Starved | -0.214 | 0.295 | 292 | -0.725 | 0.8870 |
| **Mated Starved - Virgin Starved** | **1.615** | **0.295** | **292** | **5.476** | **<.0001** |
| *(B) Pseudomonas entomophila* |  |  |  |  |  |
| **Virgin Fed - Mated Fed** | **-2.087** | **0.368** | **292** | **-5.676** | **<.0001** |
| **Virgin Fed - Mated Starved** | **-2.568** | **0.368** | **292** | **-6.984** | **<.0001** |
| **Virgin Fed - Virgin Starved** | **-2.362** | **0.368** | **292** | **-6.423** | **<.0001** |
| Mated Fed - Mated Starved | -0.481 | 0.368 | 292 | -1.308 | 0.5587 |
| Mated Fed - Virgin Starved | -0.275 | 0.368 | 292 | -0.747 | 0.8779 |
| Mated Starved - Virgin Starved | 0.206 | 0.368 | 292 | 0.561 | 0.9435 |
| *(C) Enterococcus faecalis* |  |  |  |  |  |
| **Virgin Fed - Mated Fed** | **-1.277** | **0.329** | **292** | **-3.877** | **0.0007** |
| **Virgin Fed - Mated Starved** | **-1.493** | **0.329** | **292** | **-4.534** | **<.0001** |
| **Virgin Fed - Virgin Starved** | **-2.278** | **0.329** | **292** | **-6.916** | **<.0001** |
| Mated Fed - Mated Starved | -0.216 | 0.329 | 292 | -0.657 | 0.9131 |
| **Mated Fed - Virgin Starved** | **-1.001** | **0.329** | **292** | **-3.038** | **0.0138** |
| Mated Starved - Virgin Starved | -0.784 | 0.329 | 292 | -2.382 | 0.0829 |

**Supplementary table S3.** Post-hoc analysis using Tukey’s HSD for pairwise comparisons of the effect of ‘treatments × HPI’ interaction on systemic bacterial load in infected females.

| Pairwise comparison | Estimate | SE | DF | t-ratio | p-value |
| --- | --- | --- | --- | --- | --- |
| *(A) Providencia rettgeri* |  |  |  |  |  |
| Virgin Fed 4 - Mated Fed 4 | -1.236 | 0.417 | 292 | -2.963 | 0.0644 |
| **Virgin Fed 4 - Mated Starved 4** | **-2.144** | **0.417** | **292** | **-5.140** | **<.0001** |
| Virgin Fed 4 - Virgin Starved 4 | -0.864 | 0.417 | 292 | -2.072 | 0.4355 |
| **Virgin Fed 4 - Virgin Fed 10** | **-4.933** | **0.417** | **292** | **-11.825** | **<.0001** |
| **Virgin Fed 4 - Mated Fed 10** | **-6.305** | **0.417** | **292** | **-15.114** | **<.0001** |
| **Virgin Fed 4 - Mated Starved 10** | **-9.055** | **0.417** | **292** | **-21.707** | **<.0001** |
| **Virgin Fed 4 - Virgin Starved 10** | **-7.104** | **0.417** | **292** | **-17.030** | **<.0001** |
| Mated Fed 4 - Mated Starved 4 | -0.908 | 0.417 | 292 | -2.177 | 0.3688 |
| Mated Fed 4 - Virgin Starved 4 | 0.372 | 0.417 | 292 | 0.891 | 0.9867 |
| **Mated Fed 4 - Virgin Fed 10** | **-3.697** | **0.417** | **292** | **-8.862** | **<.0001** |
| **Mated Fed 4 - Mated Fed 10** | **-5.069** | **0.417** | **292** | **-12.151** | **<.0001** |
| **Mated Fed 4 - Mated Starved 10** | **-7.819** | **0.417** | **292** | **-18.744** | **<.0001** |
| **Mated Fed 4 - Virgin Starved 10** | **-5.868** | **0.417** | **292** | **-14.067** | **<.0001** |
| **Mated Starved 4 - Virgin Starved 4** | **1.280** | **0.417** | **292** | **3.068** | **0.0479** |
| **Mated Starved 4 - Virgin Fed 10** | **-2.789** | **0.417** | **292** | **-6.685** | **<.0001** |
| **Mated Starved 4 - Mated Fed 10** | **-4.161** | **0.417** | **292** | **-9.974** | **<.0001** |
| **Mated Starved 4 - Mated Starved 10** | **-6.911** | **0.417** | **292** | **-16.567** | **<.0001** |
| **Mated Starved 4 - Virgin Starved 10** | **-4.960** | **0.417** | **292** | **-11.890** | **<.0001** |
| **Virgin Starved 4 - Virgin Fed 10** | **-4.068** | **0.417** | **292** | **-9.753** | **<.0001** |
| **Virgin Starved 4 - Mated Fed 10** | **-5.440** | **0.417** | **292** | **-13.042** | **<.0001** |
| **Virgin Starved 4 - Mated Starved 10** | **-8.191** | **0.417** | **292** | **-19.635** | **<.0001** |
| **Virgin Starved 4 - Virgin Starved 10** | **-6.240** | **0.417** | **292** | **-14.958** | **<.0001** |
| **Virgin Fed 10 - Mated Fed 10** | **-1.372** | **0.417** | **292** | **-3.289** | **0.0247** |
| **Virgin Fed 10 - Mated Starved 10** | **-4.122** | **0.417** | **292** | **-9.882** | **<.0001** |
| **Virgin Fed 10 - Virgin Starved 10** | **-2.171** | **0.417** | **292** | **-5.205** | **<.0001** |
| **Mated Fed 10 - Mated Starved 10** | **-2.750** | **0.417** | **292** | **-6.593** | **<.0001** |
| Mated Fed 10 - Virgin Starved 10 | -0.799 | 0.417 | 292 | -1.916 | 0.5408 |
| **Mated Starved 10 - Virgin Starved 10** | **1.951** | **0.417** | **292** | **4.677** | **0.0001** |
| *(B) Pseudomonas entomophila* |  |  |  |  |  |
| Virgin Fed 4 - Mated Fed 4 | -0.6918 | 0.52 | 292 | -1.330 | 0.8867 |
| **Virgin Fed 4 - Mated Starved 4** | **-1.9937** | **0.52** | **292** | **-3.834** | **0.0038** |
| Virgin Fed 4 - Virgin Starved 4 | -1.5357 | 0.52 | 292 | -2.953 | 0.0661 |
| **Virgin Fed 4 - Virgin Fed 10** | **-2.2905** | **0.52** | **292** | **-4.404** | **0.0004** |
| **Virgin Fed 4 - Mated Fed 10** | **-5.7733** | **0.52** | **292** | **-11.101** | **<.0001** |
| **Virgin Fed 4 - Mated Starved 10** | **-5.4333** | **0.52** | **292** | **-10.448** | **<.0001** |
| **Virgin Fed 4 - Virgin Starved 10** | **-5.4789** | **0.52** | **292** | **-10.535** | **<.0001** |
| Mated Fed 4 - Mated Starved 4 | -1.3019 | 0.52 | 292 | -2.503 | 0.1981 |
| Mated Fed 4 - Virgin Starved 4 | -0.8439 | 0.52 | 292 | -1.623 | 0.7363 |
| **Mated Fed 4 - Virgin Fed 10** | **-1.5987** | **0.52** | **292** | **-3.074** | **0.0471** |
| **Mated Fed 4 - Mated Fed 10** | **-5.0815** | **0.52** | **292** | **-9.771** | **<.0001** |
| **Mated Fed 4 - Mated Starved 10** | **-4.7415** | **0.52** | **292** | **-9.117** | **<.0001** |
| **Mated Fed 4 - Virgin Starved 10** | **-4.7871** | **0.52** | **292** | **-9.205** | **<.0001** |
| Mated Starved 4 - Virgin Starved 4 | 0.4580 | 0.52 | 292 | 0.881 | 0.9876 |
| Mated Starved 4 - Virgin Fed 10 | -0.2968 | 0.52 | 292 | -0.571 | 0.9992 |
| **Mated Starved 4 - Mated Fed 10** | **-3.7797** | **0.52** | **292** | **-7.268** | **<.0001** |
| **Mated Starved 4 - Mated Starved 10** | **-3.4396** | **0.52** | **292** | **-6.614** | **<.0001** |
| **Mated Starved 4 - Virgin Starved 10** | **-3.4852** | **0.52** | **292** | **-6.702** | **<.0001** |
| Virgin Starved 4 - Virgin Fed 10 | -0.7548 | 0.52 | 292 | -1.451 | 0.8322 |
| **Virgin Starved 4 - Mated Fed 10** | **-4.2377** | **0.52** | **292** | **-8.148** | **<.0001** |
| **Virgin Starved 4 - Mated Starved 10** | **-3.8976** | **0.52** | **292** | **-7.495** | **<.0001** |
| **Virgin Starved 4 - Virgin Starved 10** | **-3.9432** | **0.52** | **292** | **-7.582** | **<.0001** |
| **Virgin Fed 10 - Mated Fed 10** | **-3.4828** | **0.52** | **292** | **-6.697** | **<.0001** |
| **Virgin Fed 10 - Mated Starved 10** | **-3.1428** | **0.52** | **292** | **-6.043** | **<.0001** |
| **Virgin Fed 10 - Virgin Starved 10** | **-3.1884** | **0.52** | **292** | **-6.131** | **<.0001** |
| Mated Fed 10 - Mated Starved 10 | 0.3400 | 0.52 | 292 | 0.654 | 0.9980 |
| Mated Fed 10 - Virgin Starved 10 | 0.2944 | 0.52 | 292 | 0.566 | 0.9992 |
| Mated Starved 10 - Virgin Starved 10 | -0.0456 | 0.52 | 292 | -0.088 | 1.0000 |
| *(C) Enterococcus faecalis* |  |  |  |  |  |
| Virgin Fed 4 - Mated Fed 4 | -0.991 | 0.466 | 292 | -2.128 | 0.3994 |
| Virgin Fed 4 - Mated Starved 4 | -0.613 | 0.466 | 292 | -1.315 | 0.8926 |
| Virgin Fed 4 - Virgin Starved 4 | -1.413 | 0.466 | 292 | -3.033 | 0.0530 |
| **Virgin Fed 4 - Virgin Fed 10** | **-1.939** | **0.466** | **292** | **-4.163** | **0.0011** |
| **Virgin Fed 4 - Mated Fed 10** | **-3.502** | **0.466** | **292** | **-7.518** | **<.0001** |
| **Virgin Fed 4 - Mated Starved 10** | **-4.313** | **0.466** | **292** | **-9.260** | **<.0001** |
| **Virgin Fed 4 - Virgin Starved 10** | **-5.082** | **0.466** | **292** | **-10.910** | **<.0001** |
| Mated Fed 4 - Mated Starved 4 | 0.379 | 0.466 | 292 | 0.813 | 0.9923 |
| Mated Fed 4 - Virgin Starved 4 | -0.421 | 0.466 | 292 | -0.905 | 0.9855 |
| Mated Fed 4 - Virgin Fed 10 | -0.948 | 0.466 | 292 | -2.035 | 0.4603 |
| **Mated Fed 4 - Mated Fed 10** | **-2.511** | **0.466** | **292** | **-5.390** | **<.0001** |
| **Mated Fed 4 - Mated Starved 10** | **-3.322** | **0.466** | **292** | **-7.132** | **<.0001** |
| **Mated Fed 4 - Virgin Starved 10** | **-4.091** | **0.466** | **292** | **-8.782** | **<.0001** |
| Mated Starved 4 - Virgin Starved 4 | -0.800 | 0.466 | 292 | -1.718 | 0.6757 |
| Mated Starved 4 - Virgin Fed 10 | -1.326 | 0.466 | 292 | -2.848 | 0.0876 |
| **Mated Starved 4 - Mated Fed 10** | **-2.889** | **0.466** | **292** | **-6.203** | **<.0001** |
| **Mated Starved 4 - Mated Starved 10** | **-3.701** | **0.466** | **292** | **-7.945** | **<.0001** |
| **Mated Starved 4 - Virgin Starved 10** | **-4.469** | **0.466** | **292** | **-9.595** | **<.0001** |
| Virgin Starved 4 - Virgin Fed 10 | -0.526 | 0.466 | 292 | -1.130 | 0.9498 |
| **Virgin Starved 4 - Mated Fed 10** | **-2.089** | **0.466** | **292** | **-4.485** | **0.0003** |
| **Virgin Starved 4 - Mated Starved 10** | **-2.901** | **0.466** | **292** | **-6.227** | **<.0001** |
| **Virgin Starved 4 - Virgin Starved 10** | **-3.669** | **0.466** | **292** | **-7.877** | **<.0001** |
| **Virgin Fed 10 - Mated Fed 10** | **-1.563** | **0.466** | **292** | **-3.355** | **0.0200** |
| **Virgin Fed 10 - Mated Starved 10** | **-2.374** | **0.466** | **292** | **-5.097** | **<.0001** |
| **Virgin Fed 10 - Virgin Starved 10** | **-3.143** | **0.466** | **292** | **-6.747** | **<.0001** |
| Mated Fed 10 - Mated Starved 10 | -0.811 | 0.466 | 292 | -1.742 | 0.6600 |
| **Mated Fed 10 - Virgin Starved 10** | **-1.580** | **0.466** | **292** | **-3.392** | **0.0178** |
| Mated Starved 10 - Virgin Starved 10 | -0.769 | 0.466 | 292 | -1.650 | 0.7191 |

